# Hippocampal spike-time correlations and place-field overlaps during open field foraging

**DOI:** 10.1101/801209

**Authors:** Mauro M. Monsalve-Mercado, Yasser Roudi

## Abstract

Phase precessing place cells encode spatial information on fine timescales via the timing of their spikes. This *phase code* has been extensively studied on linear tracks and for short runs in the open field. However, less is known about the phase code on unconstrained trajectories lasting tens of minutes, typical of open field foraging. In previous work (Monsalve-Mercado and Leibold, 2017), an analytic expression was derived for the spike-time cross-correlation between phase precessing place cells during natural foraging in the open field. This expression makes two predictions on how this phase code differs from the linear track case: cross-correlations are symmetric with respect to time, and they represent the distance between pairs of place fields in that the theta-filtered cross-correlations around zero time-lag are positive for cells with nearby fields while they are negative for those with fields further apart. Here we analyze several available open field recordings and show that these predictions hold for pairs of CA1 place cells. We also show that the relationship remains during remapping in CA1, and it is also present in place cells in area CA3. For CA1 place cells of Fmr1-null mice, which exhibit normal place fields but somewhat weaker temporal coordination with respect to theta compared to wild type, the cross-correlations still remain symmetric but the relationship to place field overlap is largely lost. The relationship discussed here describes how spatial information is communicated by place cells to downstream areas in a finer theta-timescale, relevant for learning and memory formation in behavioural tasks lasting tens of minutes in the open field.

## 1 Introduction

A place cell in the hippocampus exhibits higher firing rates during traversals of specific regions of the environment, namely its place field (O’Keefe and Dostrovsky, 1971). Together as a population, these place cells form a neural representation of space from which the position of the animal can be accurately decoded (Wilson and McNaughton, 1993). In addition to this *rate code*, place cells convey spatial information at a finer time scale through the relative timing of their action potentials: the spiking activity of the cell population is organized with respect to an ongoing oscillation in the extracellular potential within the theta range (4-12 Hz) (John O’Keefe and Recce, 1993; Skaggs et al., 1996; Schmidt et al., 2009; Colgin, 2013). More precisely, a cell starts firing at a late phase of the reference theta oscillation when the animal enters the corresponding place field, progressively moving to earlier phases in each new theta cycle as the animal traverses the entire place field. The phase of the spike relative to the ongoing theta oscillation thus signals the position of the animal within the place field.

In a 1D track, phase precession causes the spikes of neurons within a theta cycle to be ordered in the same order as the place fields would be visited (Skaggs et al., 1996), thus providing a time compressed representation of place cell activity (Dragoi and Buzsáki, 2006; Melamed et al., 2004); see also Fig. 1 in Jaramillo and Kempter, 2017 and Fig. 2 of Lenck-Santini and Holmes, 2008 for schematic drawings explaining this phenomena. In fact, in recordings on a linear track, phase precession of two cells with overlapping fields can be used to directly read out how far apart the fields are from each other. This is because a cell firing early in the theta cycle signals the end of its place field and a cell firing late in the cycle signals the start, and therefore, spikes from the cells whose place fields come near the end happen earlier than the ones from a cell whose field is just being entered. The distance between two place field centers is thus proportional to the average theta phase difference of the respective cells among the several cycles where the fields overlap (Schmidt et al., 2009; Feng et al., 2015). Equivalently, the average phase difference corresponds to the time lag of the nearest peak to zero in the cross-correlation of their spike times within theta time scales. Phase precession then encodes spatial relational information between place cells, evident only over the timescale of seconds that takes for the animal to get from one field to another, into a temporal code available in the scale of tens of milliseconds, namely in the temporal order of the spikes that the cells emit. Since plasticity mechanisms such as spike-timing-dependent plasticity operate at milliseconds timescale, this encoding of the distances in the milliseconds timescale may play a major role in learning and memory consolidation (Gerstner et al., 1996; Markram et al., 1997; Bi et al., 1998; Melamed et al., 2004).

**Figure 1:**
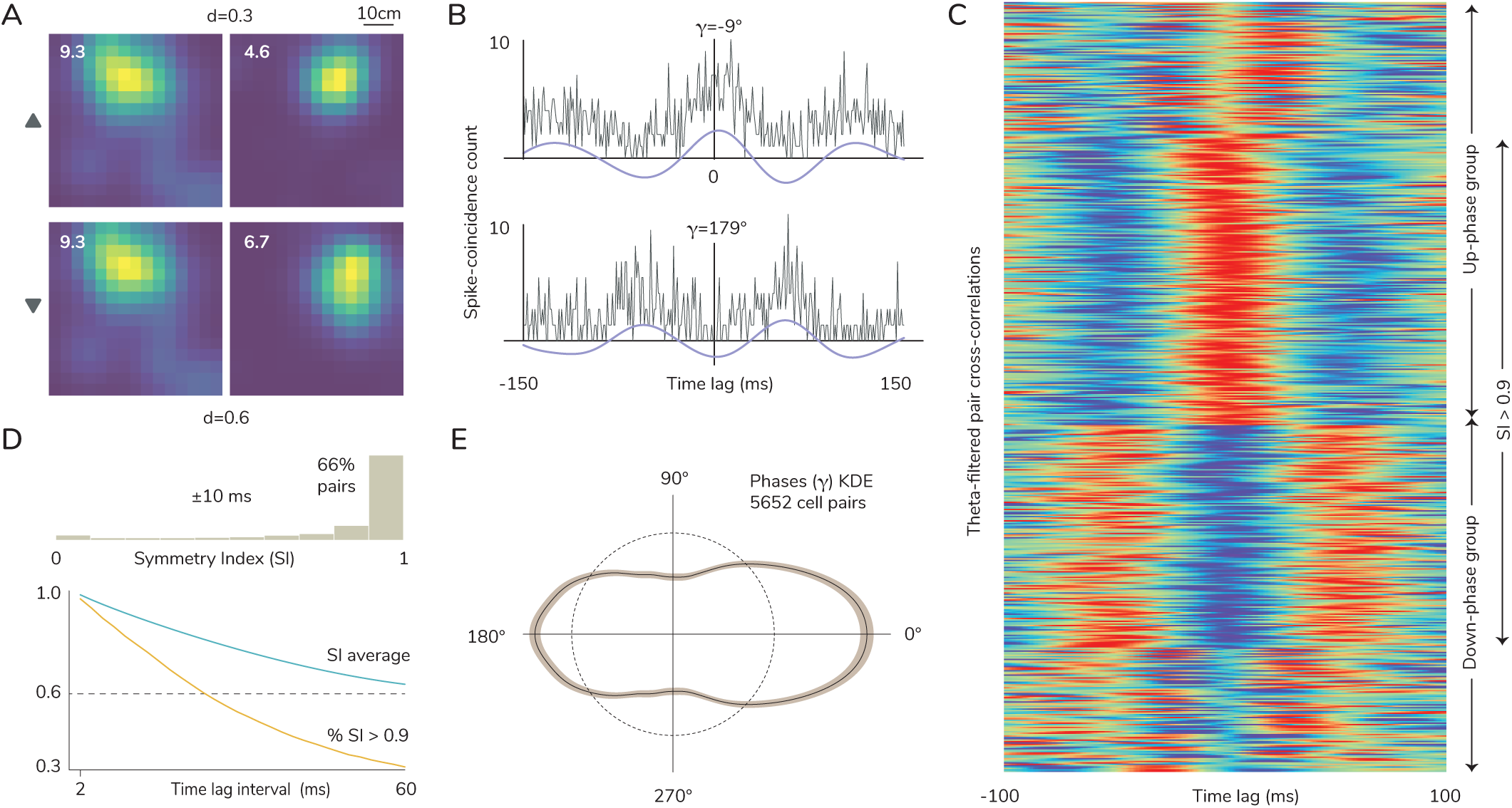
Spike-time cross-correlations are symmetric in the theta band. (A) An example illustrating the theoretical predictions. Two simultaneously recorded cell pairs are shown to have place fields close together (top row, up-triangle mark) and further apart (bottom row, down-triangle mark). The peak firing rate is shown on the corner of each rate map recorded on a 60 cm square enclosure. The Kolmogorov-Smirnov distance d between the distributions of spike locations for each pair is shown above and below their firing rate maps. (B) The corresponding cross-correlations of their spike trains are shown to the right of each pair. The theory predicts that for the top pair, whose place fields are close together, the instantaneous phase γ at zero lag of the theta-filtered cross-correlation (blue trace) is close to 0°. For the bottom pair, whose place fields are further apart but substantially overlapping, the phase is close to 180°. (C) The normalized theta-filtered cross-correlations (blue traces in A) for all 5652 cell pairs present a high degree of symmetry on theta timescales. Both the up- and down-phase groups (phases in the intervals from −90° to 90° and 90° to −90° counter-clockwise respectively) independently present a high proportion of cell pairs with a symmetry index higher than 0.9 (68% and 64% respectively). (D) The symmetry index (SI) measures the degree of symmetry of the theta-filtered cross-correlations in an interval around zero lag. The histogram of SI values (top) shows that most cell pairs have highly symmetric correlations on an interval of 20 ms centered at zero lag, with 66% of cell pairs belonging to the upper 10% (SI > 0.9) range of SI values. A great majority of cell pairs retain a high degree of symmetry in their correlations for time lag intervals relevant for synaptic plasticity (bottom: the green trace is the average of SI values for all cell pairs, and the yellow trace is the fraction of cell pairs with a SI higher than 0.9). (E) The circular kernel density estimation of the distribution of phases shows a distinctive tendency towards a bimodal distribution oriented on the 0 *−* 180° axis, although it contains a higher proportion of phases (55%) in the up-phase group.

**Figure 2:**
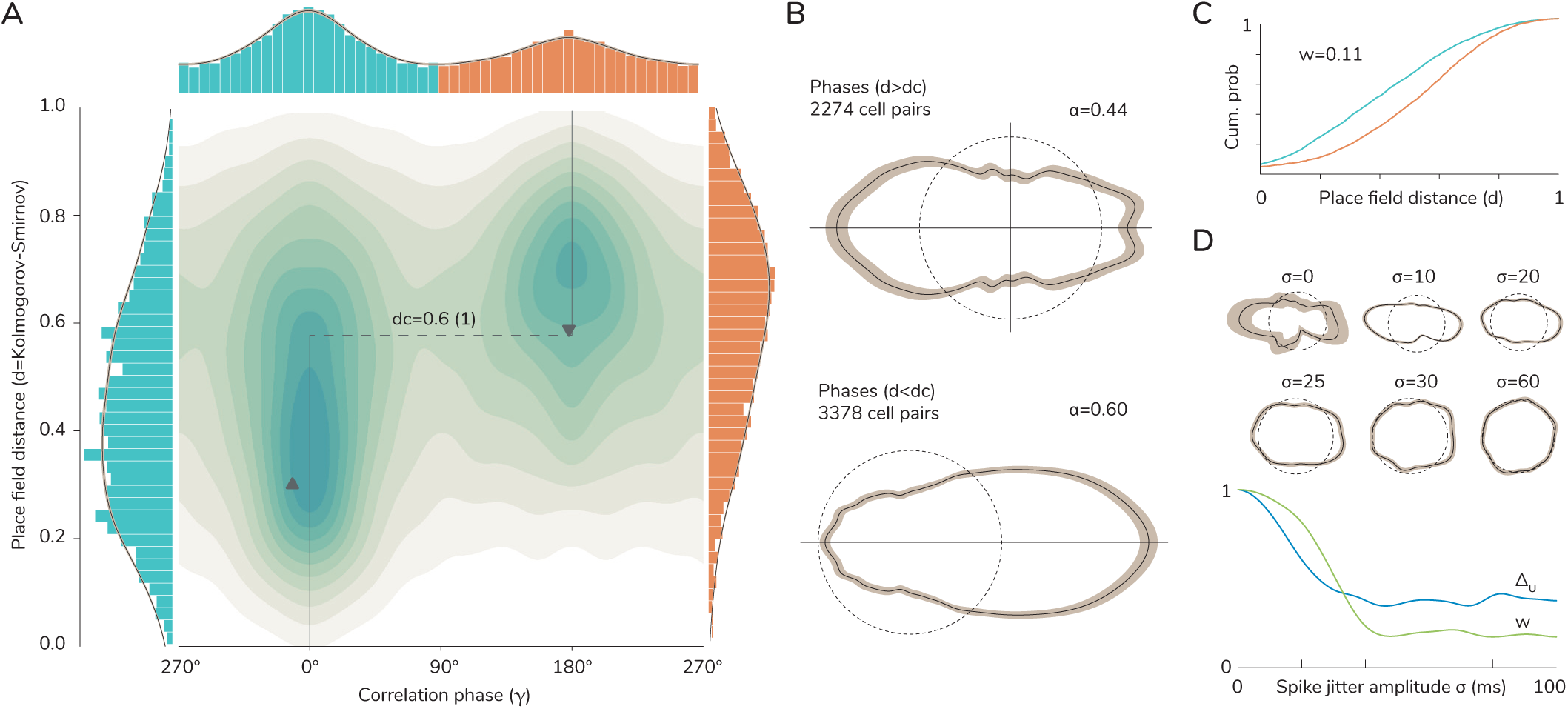
The binary phase of the correlation tells apart high-from low-overlapping fields. (A) The circular-linear kernel density estimation (CLKDE) for the joint distribution of phases and distances shows a clear separation of the data into two distinct groups in phase-distance space. The two triangular marks denote the two cell pair examples shown in Fig. 1A. The theory predicts the data should follow a step function with values at 0° and 180°. The fit minimizing circular distance shows the data is best separated at a critical distance of *d_c_* = 0.6(1). The marginal distribution of phases (on top) is conceptually divided into up-phases (blue) for the interval between *−*90° and 90° counter-clockwise and into down-phases (orange) otherwise. The marginal distributions of distances restricted to the up- and down-phase groups are shown to the left (blue) and right (orange) respectively. (B) The marginal distributions of phases restricted to distances above (top) and below (bottom) the critical distance show the majority of phases belonging to the down- and up-phase groups respectively. (C) The cumulative probability functions of the marginal distributions of place field distances reveal differences on their horizontal concentration of mass, measured by the total area between the curves (Wasserstein distance *w*). (D) The introduction of Gaussian spike jitter to a single recording session reveals that the phase-distance relationship is robust for small jitter amplitudes but it breaks down for higher amplitudes falling within theta timescales that still leave place field distances invariant. The jitter amplitude is the standard deviation *σ* of Gaussian noise. Phase distributions for different noise levels (top circular plots, made with 30 iterations of noise each) suggest a tendency for phases to become uniformly distributed with increasing noise, which is confirmed by their decreasing circular Wasserstein distances from uniformity Δ*_U_* (blue trace). Even though place field distances remain invariant, they become disorganized with respect to their associated phase values, resulting in decreasing *w* with noise and the loss of the phase-distance relationship. The values of *w* and Δ*_U_* are normalized to the case without jitter.

Although initially discovered in 1D environments, phase precession was later shown to be present also in open field two-dimensional recordings (Skaggs et al., 1996; Jeewajee et al., 2014; Huxter et al., 2008). Phase precession in these studies is similar to the recordings on linear tracks. For instance, as in the 1D case, spikes precess over the whole range of the theta cycle (approximately 360°) also in 2D (Jeewajee et al., 2014). In addition, the firing phase of a cell correlates with different measures of the distance traversed within the field. And more importantly, for cells with overlapping fields, the shift in the peak of their cross-correlation negatively correlates with the amount of field overlap, although this depends on the direction of running, it is weaker for runs near the periphery of the fields and for runs at low speed (Huxter et al., 2008).

To analyze phase precession in open field trajectories, past work often focuses on seconds-long high-speed short segments of the whole trajectory as independent single runs through a cell’s place field, usually defined from the moment of entry to the field until the next exit. However, natural foraging behaviour in open field environments usually involves exploration in the order of tens of minutes and on unconstrained trajectories away from stereotypical straight runs. Spatial learning and memory tasks in the open field typically require several learning trials, lasting several minutes each, to obtain satisfactory performance levels (Vorhees and Williams, 2014). Similarly, although place cells usually form rapidly upon exploring a novel environment (Hill, 1978), it takes several minutes of exposure to reach a new independent and stable place field code, over which slowly developing plastic changes improve spatial tuning, field stability over trials, and spatial information content (Frank et al., 2004; Wilson and McNaughton, 1993). All these warrant a better understanding of the properties of the phase code during natural foraging.

In previous theoretical work (Monsalve-Mercado and Leibold, 2017), we make predictions about the correlations between phase precessing place cells during natural foraging of open field environments and how these correlations may encode spatial information. This is done by modelling a single cell’s firing pattern as a Gaussian spatial envelope modulated by an oscillatory component with a frequency slightly higher than theta, and averaging the correlation between such pairs of place cells over all straight paths crossing the area of overlap between the place fields. The analysis shows that place cells with high field overlap present no average phase difference with respect to theta, while cells with less but substantially overlapping fields have an average half a cycle phase difference. In other words, minutes-long cross-correlations filtered at theta frequency are positive around zero time-lag for cells with nearby fields while they are negative for those with fields further apart. In the following we refer to this effect as the *phase-distance* relationship. In the same study, it was shown that this relationship could be instrumental in the emergence of grid fields in the entorhinal cortex. Briefly, this phase-distance relationship in the cross-correlations, combined with spike-timing-dependent plasticity in projections from place cells to a model grid cell, was shown to induce effective Mexican-hat like interactions between the projections from place cells which then leads to a hexagonal pattern.

The aim of the current paper is to see whether the main theoretical predictions about the pair-wise cross-correlations, in particular their relationship to the spatial overlap between the fields holds in real data during natural foraging. To do this, we analyzed several electrophysiological recordings of hippocampal place cells. We found that the qualitative features predicted about the cross-correlation holds in the CA1 datasets that we analyzed. The relationship also persists during remapping experiments in CA1 as well as in data analyzed from CA3. To further look into the relationship between the phase-distance effect and the timing of spikes with respect to theta, we also analyzed data from Fmr1-null knockout mice in which place fields appear to be normal but their activities appear to be less organized with respect to theta. We found that in these cells the cross-correlations remain symmetric, but the relationship between the cross-correlations and the spatial overlap of place fields is largely lost.

## 2 Methods

### 2.1 Predictions of the model

We model the activity of a population of CA1 place cells for an animal engaged in an open field exploratory task. A place cell is strongly modulated both in space and time, firing within a localized area of the environment (its place field), and only periodically in time in accordance with phase precession. We model this as a Gaussian envelope modulated by an oscillatory component. Mathematically, this is done by writing the firing rate, *H_n_*(*t*) of neuron *n* at time *t* and when the animal is at position *x*(*t*) as

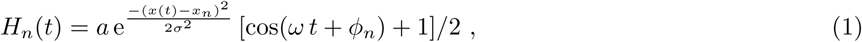

where *ω/*(2*π*) is the single neuron oscillatory frequency, *x_n_* is the centre of the place field of neuron *n*, *φ_n_* is the oscillation phase, *σ* controls the size of the place field, while *a* determines the maximum firing rate. Theta phase precession naturally arises by taking the oscillation frequency *ω/*(2*π*) in Eq. 1 to be slightly higher than the theta oscillation frequency of 8Hz.

The cross-correlation function of the activity of two cells *n* and *n′* is defined as

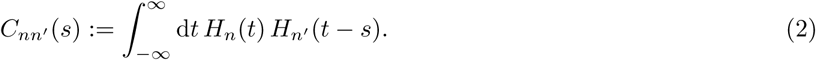

The cells’ activities contribute the most to the correlation for paths crossing a region of high place field overlap. To compute the cross-correlations, *C_nn′_*, (Monsalve-Mercado and Leibold, 2017) assumed that the time integral in Eq. 2 could be re-written as an average over straight paths of all possible orientations traversing the midpoint between the centers of the place fields of neurons *n* and *n′*. They showed that for cells with equal peak firing rate and place field size, and for constant speed and small time lags *s*, the cross-correlation can be approximated as

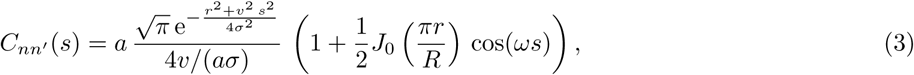

where 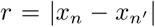 is the distance between the place field centers, *v* the speed of the animal, *R* is the distance from the place field centre at which the firing rate has decreased to 10%, and *J*_0_ is the Bessel function of the first kind.

Two important predictions can be inferred from the expression for the correlation function in Eq. 3 that are a direct consequence of the two-dimensional symmetry of open field exploration:

The correlation is symmetric in the time domain. This can be easily seen by the fact that changing *s* to *−s* in Eq. 3 does not change anything. An important implication is that the oscillatory component of the correlation around time-lag zero must be in either a peak or a valley. In other words, when filtered in the theta range, cells can be strongly positively or negatively correlated. This organization is not arbitrary, but results in the second prediction.

Distance is reflected in the phase of the correlation at time-lag zero. Cells whose place fields are close together (have high overlap) are positively correlated in the theta range (their correlation is near a peak), while cells whose fields are further away from each other (little overlap, but still significant) are negatively correlated in the theta range (their correlation is near a valley).

### 2.2 Analysis

For each recording session we computed spike train correlations between all cell pairs for the entire duration of the recording. Only cells with more than a 100 spikes were included in the analysis. Spike trains were considered isolated events with a 1 ms accuracy, for which the correlation measures the number of coincident spikes for each time lag with a resolution of 1 ms within a 300 ms range. The correlation is filtered in the theta band (5*−*12 Hz) and the analytical signal is obtained from its Hilbert transform. We identify each cell pair with the instantaneous phase *γ* of its analytical signal at zero time lag. Only cell pairs were included in the analysis for which the instantaneous envelope of its analytical signal (not the value of the oscillation) at zero time lag was above a 0.2 threshold in absolute value.

We define a measure to evaluate the symmetry of the filtered cross-correlation 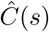. The symmetry index (SI) is defined as the normalized square integral of the symmetric part of 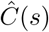 for a specific interval *±τ* around zero time lag:

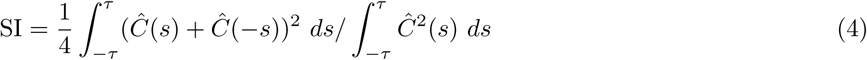

With this definition, the symmetry index, SI, ranges between 0 reflecting total antisymmetry to 1, representing total symmetry, thus quantifying the degree of symmetry in the filtered cross-correlation.

The distance *d* between two place fields was computed using the two-dimensional Kolmogorov-Smirnov probability distance directly from the distributions of spike locations (Peacock, 1983). The Wasserstein distance *w* is used to compare the marginal distance distributions, it corresponds to the area between the cumulative probability distributions. The circular Wasserstein distance Δ is used to compare the distribution of phases to another distribution with circular symmetry, it corresponds to the minimal distance from all linear Wasserstein distances on an unfolded circle for all possible starting points on the circle (Rabin et al., 2011). Circular and circular-linear kernel density estimations use von Mises and Gaussian kernels with adaptive concentration and smoothing parameters (Garćıa-Portugúes et al., 2013).

## 3 Results

We examined several published datasets of extracellular recordings in hippocampal areas CA1 and CA3 during open field exploratory foraging. The following datasets were included in the analysis: 2 CA1 sessions from a teleportation experiment reported in (Jezek et al., 2011), 5 CA1 sessions recorded in the Buzsáki lab (K. Mizuseki et al., n.d.; Kenji Mizuseki, Diba, et al., 2014; Kenji Mizuseki, Sirota, et al., 2009) (specifically session ec14.215, ec14.277, ec14.333, ec14.260, and ec15.047 from the openly available hc-3 dataset), 16 CA1 sessions from wild-type mice and 16 CA1 sessions from Fmr1-null mice recorded from (Sparks et al., 2017; Talbot et al., 2018; Dvorak et al., 2018), also taken from the openly available hc-16 dataset, 28 CA1 sessions in 3 rats from a remapping experiment reported in (Schlesiger, Cannova, et al., 2015; Schlesiger, Boublil, et al., 2018) and, finally, 178 CA3 sessions from 7 rats in 11 rooms reported in (Alme et al., 2014). Details about the recordings and the experimental settings can be found in the respective references.

Fig. 1A,B shows an example illustrating the two theoretical predictions. Fig. 1A shows the firing rate map of 3 simultaneously recorded CA1 place cells from (Jezek et al., 2011). Two possible pairs are shown in the top and bottom rows for comparison, and to the right of each row we show the respective cross-correlations of their spike times (Fig. 1B). The distributions of spike locations for the top pair are significantly closer than those of their bottom counterpart (Kolmogorov-Smirnov distance *d* = 0.3 and *d* = 0.6, respectively). The corresponding cross-correlations, averaged over approximately 10 minutes of free foraging, exhibit a high degree of symmetry around zero time lag, with symmetry indices SI = 0.95 and 0.98, respectively; see Eq. 4 for the definition of SI which ranges from 0 (total anti-symmetry) and 1 (total symmetry). In addition, the filtered correlations in the theta band (blue traces), show that the top neurons are positively correlated in theta with the phase of the peak *γ* = *−*9°, while the bottom neurons are negatively correlated with *γ* = 179°. In the following sections we explore these properties for different cells and experiments from CA1 and CA3.

### 3.1 CA1 spike-time correlations present a high degree of symmetry

At the population level, we found that 5652 cell pairs (about 47% of total pairs) passed the criteria to be included in the analysis of CA1 datasets. Overall, we observe a qualitative high degree of symmetry in the filtered cross-correlations in the theta band, even for ranges of a whole theta cycle (Fig. 1C and 1D). For each cell pair, we computed the symmetry index (SI) for intervals between 2 and 60 ms centered at zero time lag, which are relevant timescales to trigger spike-timing dependent plasticity rules (Markram et al., 1997). We observe that a majority of cell pairs present a high degree of symmetry in their correlations (Fig. 1D), especially for ranges close to zero time lag. Both the average of SI values and the fraction of cell pairs with a SI higher than 0.9 decrease when the SI is computed for increasingly larger time lag intervals, with the average SI leveling off to 0.6 for intervals up to 600 ms.

We additionally computed a circular kernel density estimation (CKDE) for the distribution of phases from all cell pairs. Since identity reversal of cell pairs is equivalent to a mirror transformation in the correlation function, to avoid ambiguity in the choice of cell-pair phase we include both possible phases for each cell pair in the computation of the CKDE. This is justified by the high degree of symmetry of a majority of correlations, and randomly choosing one of the two phases produces qualitatively similar results (not shown). As a result the CKDE displays a mirror symmetry with respect to the horizontal axis (Fig. 1E).

We found the distribution of phases to be significantly different from uniform (Hermans-Rasson, Ajne, and Watson tests: p *<* 0.001 by bootstraping with replacement). Next we wanted to characterize how the distribution of phases concentrate around 0° and 180°. For this, we computed the circular Wasserstein distance between the distribution of phases and a family of biased bimodal delta distributions oriented on the 0 *−* 180° axis, *αδ*(*γ*) + (1 *− α*)*δ*(*γ −* 180°), where the bias parameter *α*, taking values from 0 to 1, measures how strongly the distribution is inclined towards 0°. We select from the family the distribution that minimizes the distance, corresponding to a specific value of *α*. For the CA1 dataset, we obtained a bias value of *α* = 0.54, consistent with a 55% of the population of phases tending towards 0° (up-phase group), that is the proportion of phases in the interval from *−*90° to 90° counter-clockwise. The circular Wasserstein distance from the biased bimodal distribution Δ*_B_* = 0.08 is a useful measure to compare the bimodality of the distribution of phases across sessions, animals, brain regions, and experiments.

### 3.2 The binary phase of the correlation tells apart high-from low-overlapping fields

The second prediction of the model is concerned with the relationship between the phase associated to a cell pair and the corresponding distance of their place fields, i.e. a *phase-distance* relationship. As a measure of place field distance, we compute the Kolmogorov-Smirnov distance between the distributions of spike locations of both fields. This measure takes values between 0 and 1, and for a pair of two-dimensional Gaussians of equal size increases linearly with the distance between their peaks and saturates quickly after the overlap is minimal. The KS distance has the advantage of highlighting regions of substantial overlap while being a more robust measure of distance than the spatial correlation, distance between the firing rate peaks, and amount of field overlap, all of which result in qualitatively similar results at the population level (not shown).

We obtained the KS distance for all cell pairs in the CA1 datasets and computed a circular-linear kernel density estimation for the distribution of phases and distances (Fig. 2A). The CLKDE shows a qualitative preference for small and large distances to be clustered around phases of 0° and 180° respectively. To quantify how well the theoretical prediction explains the observed distribution, we fit a step function with values at 0° and 180° to determine the critical distance *d_c_* that minimizes the circular distance from the data to the step function. We found that a critical distance of *d_c_* = 0.6(1) explains best the distribution. The marginal phase distributions for small (*d < d_c_*) and large (*d > d_c_*) distances (Fig. 2B) show a preference for 0° and 180°, with bias parameters of *α* = 0.60 (59% of up-phases) and *α* = 0.44 (44% of up-phases) respectively. In addition, the marginal phase distributions show similar circular Wasserstein distances from a bimodal distribution (Δ*_B_* = 0.08 for both) as the complete CA1 dataset.

We can gain a different perspective on the separation of distances by their corresponding phases by examining the marginal distributions of distances for up- and down-phases (blue and orange distance distributions in Fig. 2A,C). We compute the Wasserstein distance between the marginal distributions of distances, which measures the area between the corresponding cumulative probability distributions (Fig. 2C). It highlights differences in the concentration of horizontal mass, setting apart distributions mostly representing different regions in the horizontal axis (distances). We found the two marginal distributions to be significantly different from each other (*p <* 0.001), with a Wasserstein distance of *w* = 0.11.

To test the robustness of the phase-distance relationship we introduce Gaussian noise with standard deviation σ into the spike times of a single recording session with 30 place cells (from Jezek et al., 2011). Cell pairs range from 212 to 100 depending on the noise level, since the theta amplitude of correlations are typically reduced with the jitter, and the threshold for inclusion of a pair is fixed. In Fig. 2D we show examples of phase distributions for different noise levels. Except for the case without jitter (*σ* = 0), each distribution is obtained by averaging 30 iterations of noise with the specified *σ* value. Because we expect the bimodality to be lost with noise, in this panel, we only show the phase distributions without symmetrization with respect to the horizontal axis. For increasing level of the jitter, *σ*, we also computed the Wasserstein distances between the marginal distributions of place field distances *w*, and the circular Wasserstein distance of phase distributions from uniformity Δ*_U_*. They are computed for *σ* between 0 and 100 ms in steps of 1 ms, and traces are smoothed afterwards with a Gaussian kernel of 5ms standard deviation. *w* and Δ*_U_* are normalized to the case where no jitter is present. The relationship is robust to spike jitter up to standard deviations of around a quarter of a theta cycle. For this range, cell pairs still present strong phase bimodality and high *w*. For higher noise amplitudes, phases tend to become more uniformly distributed as measured by decreasing Δ*_U_*. More importantly, phases become disentangled from place field distance as the difference *w* between the marginal distribution of distances drops below significant levels. For this jitter amplitudes the firing rate maps and thus place field distances remain mostly unchanged, implying that the drop in *w* is a result of phases and place field distances becoming disentangled. Since high *w* values could in principle be present for uniform phase distributions, this result suggests that bimodal phase distributions are required at the mechanistic level for the organization of place field distances into two independent clusters.

We examined several individual examples from the CA1 dataset to probe the consistency of the phase-distance relationship among datasets (Fig. 3). We found the relationship to be present in individual recording sessions. As an example we present the ec14.215 session of the hc-3 dataset, which has the most cells simultaneously recorded (60) and yielded the highest number of cell pairs (1239) (top row in Fig. 3). Data from two of the rats from the hc-13 dataset, namely ec16 (1061 cell pairs) and ec13 (865 cell pairs), were somehow outliers, which is why they were left out of the collection of CA1 datasets reported in section 3.1. They presented a strong bias in the distribution of phases towards 0°, with 72% (*α* = 0.72) and 76% (*α* = 0.75) of phases belonging to the up-phase group respectively. To see the effects more clearly in section 3.1 we therefore did not include them. However, we checked that in these two datasets the phases were still well represented by a bimodal distribution (Δ*_B_* = 0.09 and 0.10) and the phase-distance relationship was present, although weaker than the rest of the CA1 dataset (*w* = 0.06 and 0.08, *p <* 0.001). The relationship was present as well for individual sessions of the data from (Jezek et al., 2011) (middle row in Fig. 3). Pooled data over the two sessions in two independent rooms shows a clear tendency for phases to cluster around 180° (*α* = 0.47, 47% of up-phases, Δ*_B_* = 0.09). The corresponding separation of distances by their phases was strongest for the entire dataset (*w* = 0.17). In addition, we found the relationship present as well for mice, for which place cells are less spatially specific and phase precession and modulation by theta rhythms is slightly weaker than in rats (Mou et al., 2018); see the bottom row in Fig. 3. Pooled over 4 sessions in circular and rectangular arenas we found the summary statistics to be in good agreement with the other CA1 datasets (*α* = 0.64, Δ*_B_* = 0.09, *w* = 0.15).

**Figure 3:**
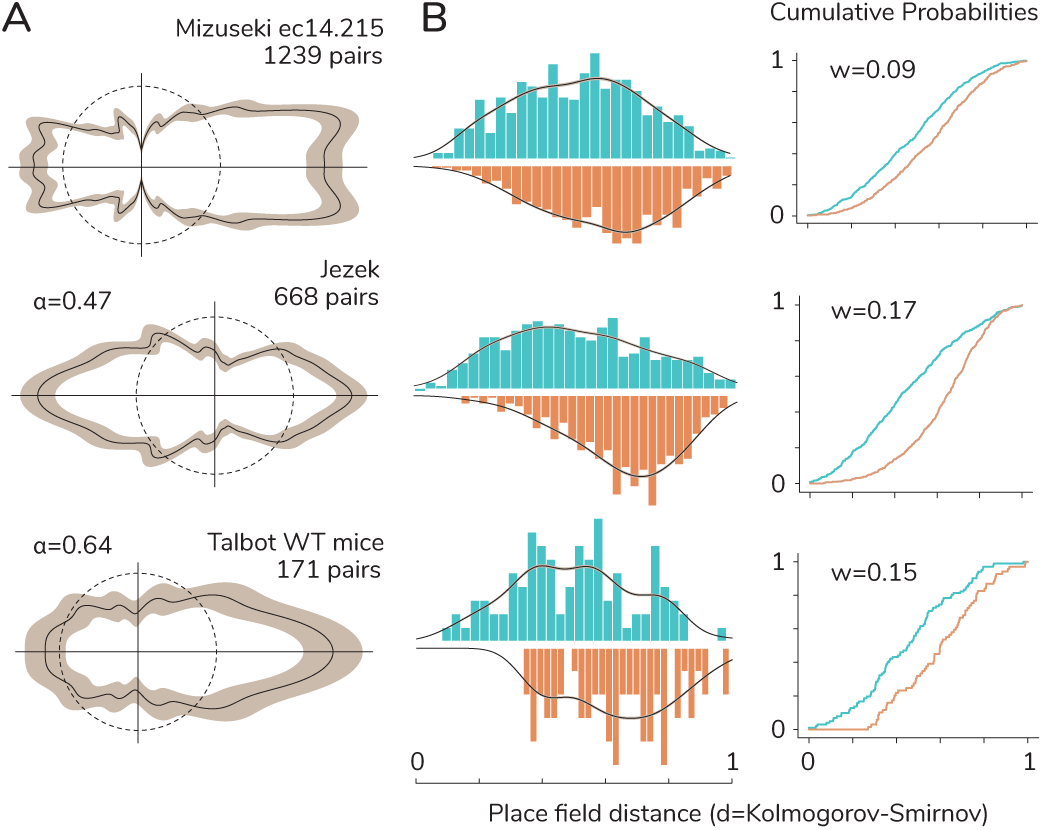
Examples from the CA1 dataset. Each row shows a different example highlighting the consistency of the phase-distance relationship. The top row shows a single recording session from animal ec14 of the hc-3 dataset. The CKDE of phases on the left panel shows clear bimodality, while middle and right panels show how the restricted marginal distance distributions differ. The middle row shows the results for the teleportation experiment for data pooled from two sessions in two independent environments and the bottom row shows the same relation present for mice as well (wild-type control mice, to compare with the results for knock-out mice in Fig. 6).

### 3.3 The phase-distance relationship is present in independent environments under global remapping

Under global remapping, place cells typically change the location of their place fields when changing environments. For two environments with independent place cell representations it is possible that cell pairs with highly overlapping fields in one environment might have low overlapping fields in the other one, for which the theory predicts a change in the phase of the cross-correlation between and up- and down-phase. We found 8 such pairs recorded from different tetrodes with a symmetry index higher than 0.8 (an example pair is shown in Fig. 4A).

**Figure 4:**
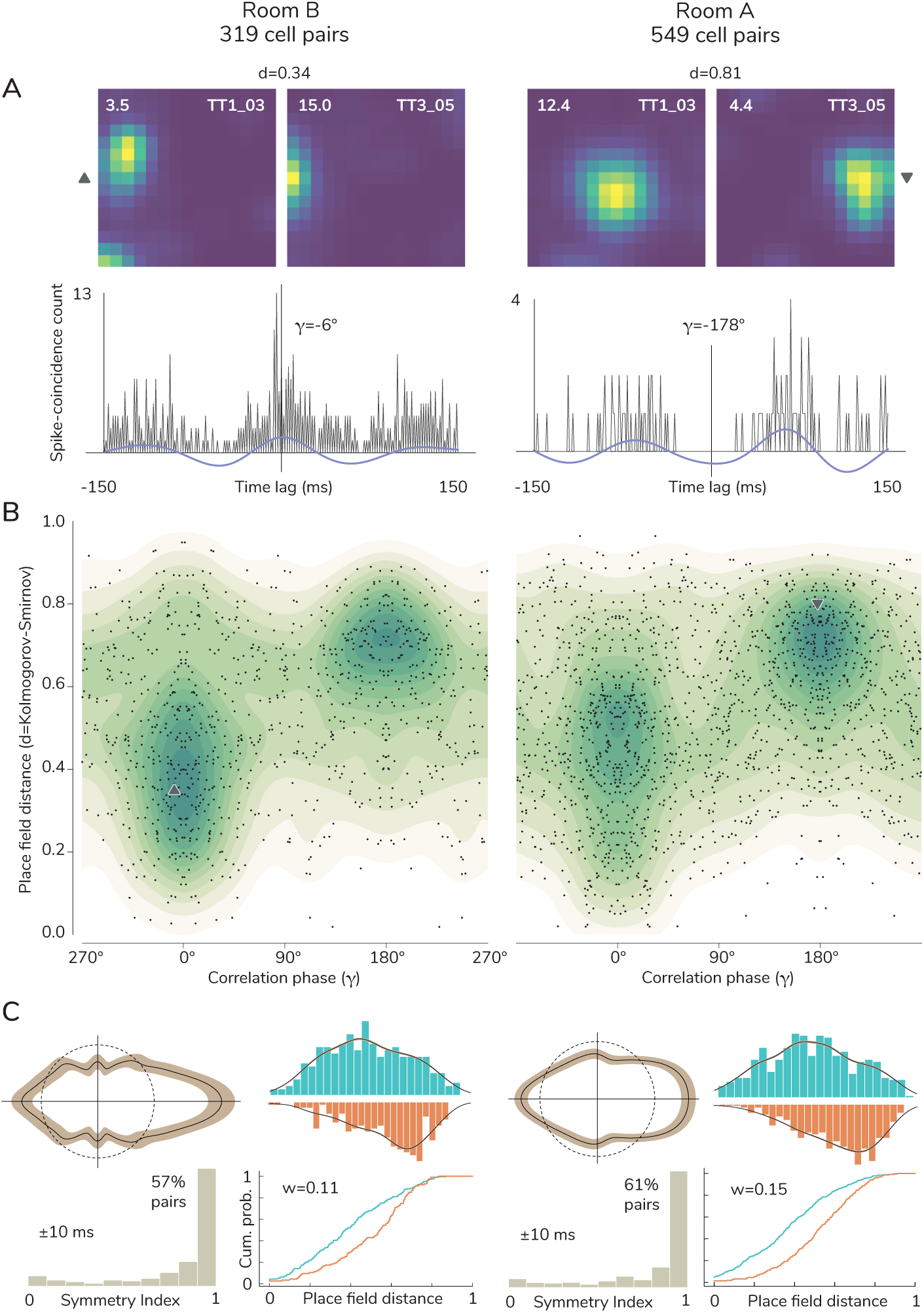
The phase-distance relation remains unchanged during remapping. (A) An example shows firing rate maps and spike-time cross-correlations for the same cell pair in two independently represented rooms (as in Fig. 1A, tetrode unit label is shown on the top right corner of the maps). It shows that a cell pair whose fields are close together and whose spiking activity is highly correlated in a given environment can change to the reverse situation in a different environment. The example illustrates the possibility for a cell pair to instantaneously change from one phase-distance subgroup to another under global remapping, while still being part of a stable phase-distance relationship at the population level (B,C). Triangular marks denote the location of each cell pair in A within their respective phase-distance distributions in panel B.

Although, most cell pairs did not show substantial overlap of their activity in both environments, we found 54 cell pairs with strong overlap and high SI (*> .*8) in both environments. Among these pairs, 42 remained in the up-phase group, and 4 remained in the down-phase group. Despite only a small number of cell pairs preserving their phase-distance relationship in both environments, the relationship was strongly present in both independently represented environments (Fig. 4B,C), although with differences in phase bias and distance separation (Room B: *α* = 0.64, 64% of up-phases, 57% of pairs with SI*>* 0.9, *w_B_* = 0.09, *w* = 0.11, Room A: *α* = 0.59, 59% of up-phases, 61% of pairs with SI*>* 0.9, *w_B_* = 0.10, *w* = 0.15). Moreover, we observe that the relationship is already in place on the first exploration of the novel environment, with identical statistics as reported for the more stable subsequent recordings in room B.

### 3.4 The phase-distance relationship is present in area CA3

Place cells in area CA3 also show phase precession with respect to the theta rhythm (John O’Keefe and Recce, 1993). We next asked whether the same phase-distance relationship exists as found in CA1. Overall, pooled data over 178 sessions from 7 rats in 11 different rooms resulted in 1121 cell pairs included in the analysis. Because the CA3 representation of space is more sparse than in CA1 (Alme et al., 2014), with only a fraction of recorded cells active in each individual room, no single session had enough cell pairs for a stand alone analysis. Only 119 out of the 178 sessions had at least two active cells with more than 100 spikes each to estimate meaningful correlations. The number of cells in each session ranged from 2 up to 17 cells, with an average number of 8.3 cells per session. Likewise, the number of cell pairs passing the selection criteria ranged from 1 up to 34, with an average of 9.8 pairs per session. We therefore had to resort to pooling the data over multiple sessions to perform our statistical analysis. Over pooled data, we found the distribution of phases to be significantly different from uniform (*p <* 0.001, Fig. 5A,B). More importantly, it was strongly biased towards 0°, with 68% of phases belonging to the up-phase group (*α* = 0.69), but still reproducing a bimodal distribution (Δ*_B_* = 0.10) with a high degree of symmetry (72% of pairs with a SI *> .*9). However, unlike most datasets in area CA1 (see the comment on rats ec16 and ec13 from the hc-3 dataset in section 3.2), the distribution of down-phases could not be concluded to be significantly different from uniform (*p* = 0.5, Kolmogorov-Smirnov test), while the up-phase group showed pronounced differences (*p <* 0.001). Despite marked differences with the CA1 dataset, the phase-distance relationship was robustly present in area CA3 (Fig. 5A,C). The marginal distance distributions were significantly different from each other (*p <* 0.001, by bootstraping with replacement), and were similarly distributed as in the CA1 region (*w* = 0.11, Fig. 5C).

**Figure 5:**
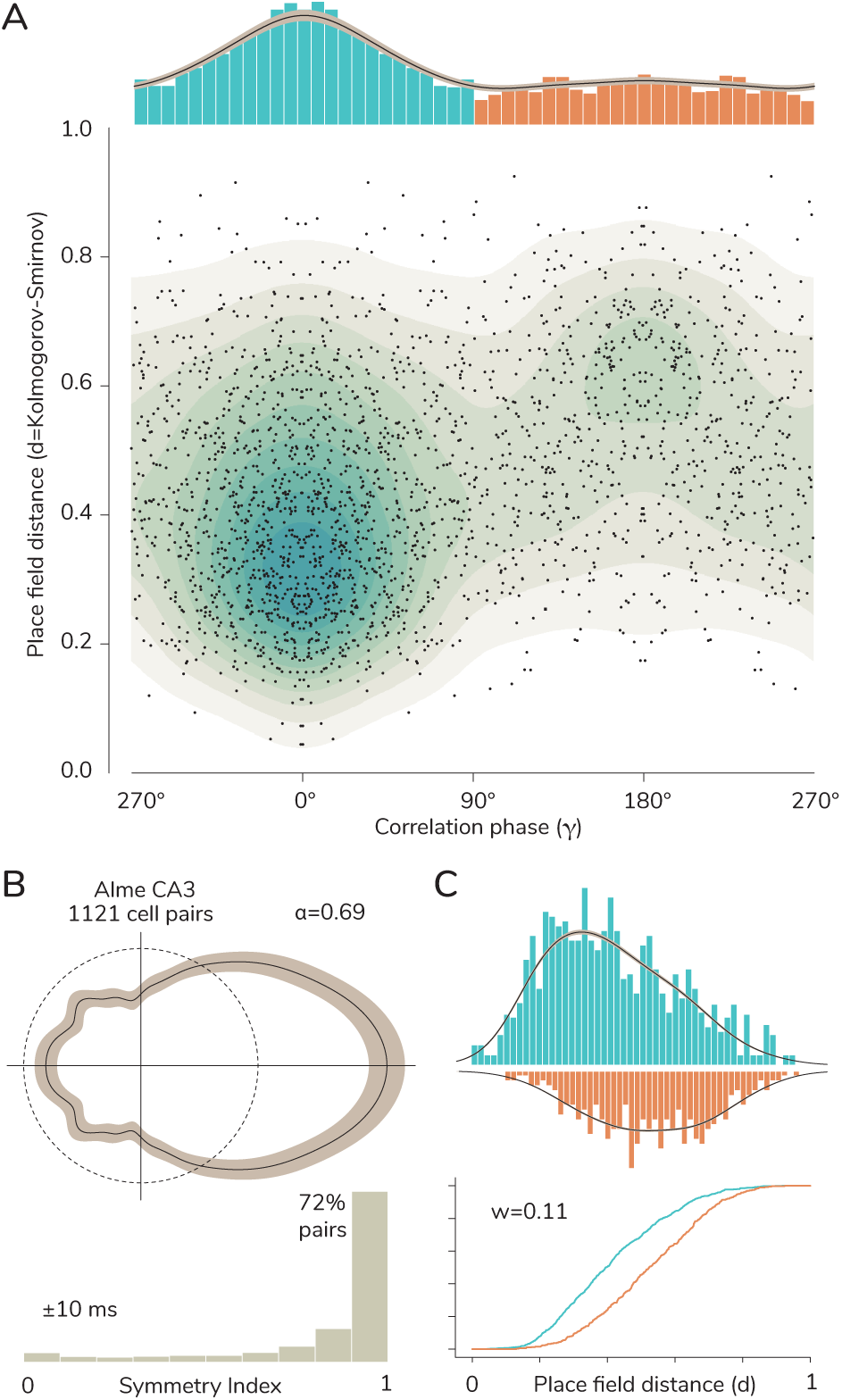
CA3 cells show the phase-distance relationship. The phase-distance CLKDE (A) shows a clear separation of the data into two subgroups, but it differs from area CA1 in that phases are highly clustered around 0° and the distribution of the down-phase group is close to uniform (B) when pooled over 178 sessions. Yet, the restricted marginal distance distributions show clear differences (C).

### 3.5 The phase-distance relationship is impaired in Fmr1-null mice

Fmr1-null mice are used as a model for fragile X syndrome, exhibiting symptoms of intellectual disability such as cognitive inflexibility. (Talbot et al., 2018) reports that, although these mice place fields are no different than wild-type, the timing of CA1 spikes are somewhat less organized by the phase of theta in the knockout mice than the wild type and also the phase-frequency discharge probabilities of the place cells appear to be less coordinated in the knockout mice than the wild type. This led us to ask whether the phase-distance relationship is disrupted in Fmr1-null mice. We found that, as in the CA3 case, the distribution of phases was strongly biased towards 0°, with 76% of phases in the up-phase group (*α* = 0.77), and was still well-represented by a bimodal distribution (Δ*_B_* = 0.09, Fig. 6A,B), with a high degree of symmetry (64% of pairs with a SI *> .*9). However, unlike in the CA3 dataset, the marginal distance distributions could not be concluded to be significantly different from each other (*p* = 0.07, by bootstraping with replacement, Fig. 6A,C). Compared to wild-type mice (bottom row in Fig. 3), the Wasserstein distance between the marginal distance distributions for Fmr1-null mice (*w* = 0.03) is much smaller than its wild-type counterpat (*w* = 0.15), indicating a diminished ability to represent disjoint regions of field overlap.

**Figure 6:**
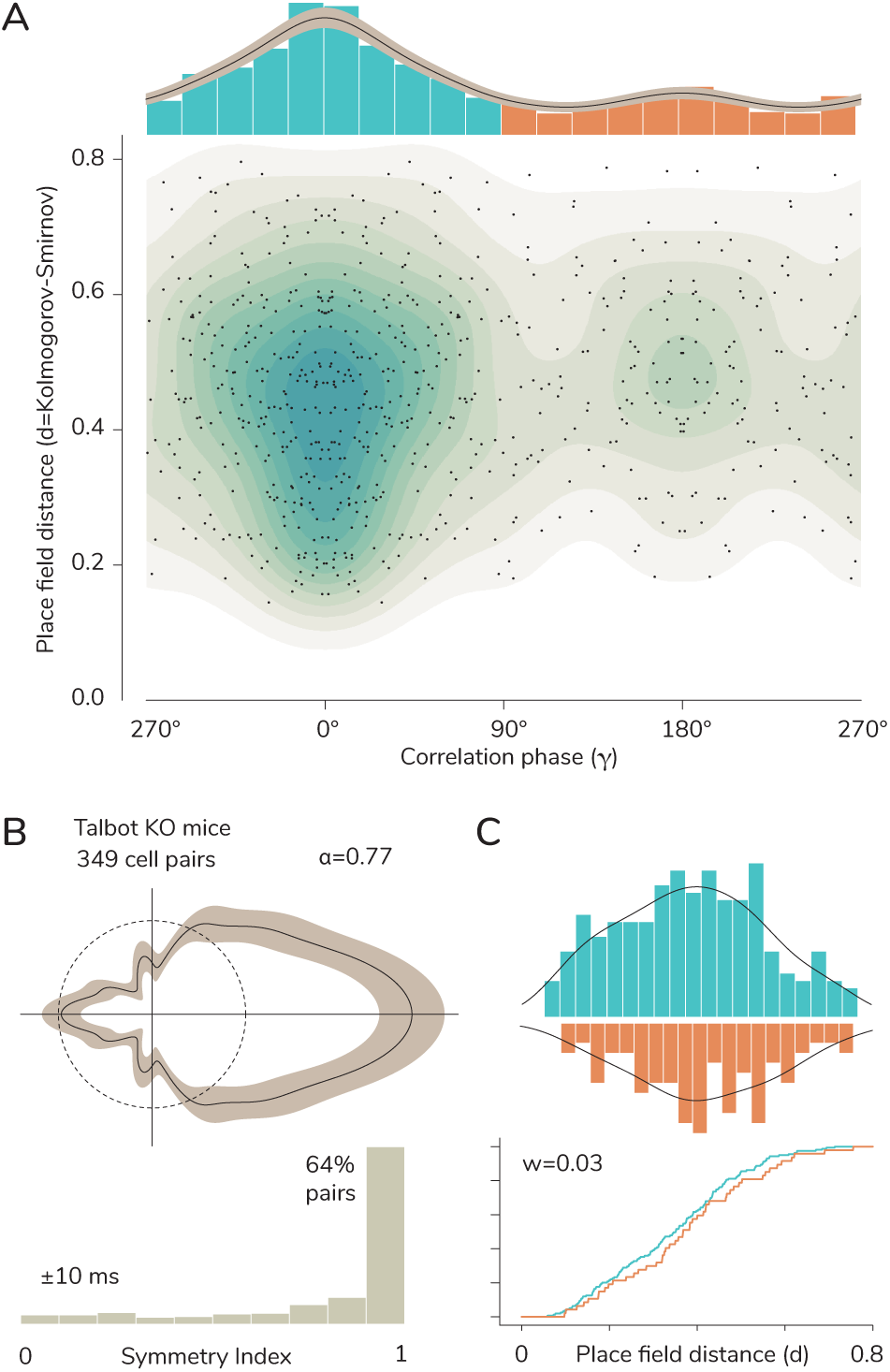
The phase-distance relationship is impaired in Fmr1-null mice. Unlike in the CA1 and CA3 areas of control animals, the phase-distance distribution for area CA1 in knock-out mice (A) shows no clear separation of the data into two subgroups simultaneously both in phase and distance, but rather only a pronounced bimodality of phases with a strong tendency towards the up-phase group (B), without significant differences in the marginal distribution of distances (C).

## 4 Discussion

The precession of hippocampal place cells’ spikes with respect to the theta rhythm is believed to be an important part of the representation of spatial information in the brain (John O’Keefe and Recce, 1993; Burgess and John O’Keefe, 1996; John O’Keefe and Burgess, 2005; Moser et al., 2008; Colgin, 2013), as it encodes the position of the animal within a field and leads to the ordering of spikes of pairs of neurons in the order the corresponding place fields are traversed (John O’Keefe and Recce, 1993; Skaggs et al., 1996).

A recent analytic estimation of the pairwise cross-correlation of place cells shows, when filtered in the theta band, that phase precession endows these cross-correlations with two properties: that they are symmetric in time-lag, and that they encode the distance between place fields in that zero time-lag is positive for cells with nearby fields while it is negative for those further apart. Analyzing short, high speed runs from one place field centre to another, Huxter et al. (Huxter et al., 2008) show results consistent with these properties, although the analytic calculations have been based on simplified assumptions such as equal size and a perfectly circular shape. In this paper, we showed that, despite these simplifying assumptions, both properties of symmetry and encoding spatial information in the cross-correlations are valid in CA1 during tens of minutes of natural foraging in the open field, in different datasets, during remapping, and even in area CA3.

The theory predicts that the phase-distance distribution should follow a step function with values at 0° and 180°. However, even though the distribution of phases is highly symmetric, we observe a high amount of variance in the circular dimension. One of the main contributors to this variance, as predicted by theoretical considerations, are biases in the animal’s trajectory, which do not fully explore every point of the environment from every possible direction in a uniform way. Besides this and other natural sources of noise, the other main source of variance is predicted to come from the spread and shape of the spatial distribution of spikes, which distort the two-dimensional symmetry assumed by the theory. An important additional mathematical prediction (unpublished), which could contribute to the observed variance, is a systematic shift in the peak of the cross-correlation of a cell pair when their fields have largely different sizes. This shift depends non-linearly on the distance between their firing rate peaks and the ratio of their field sizes, saturating at about an octave of a theta cycle for infinite distance and size ratio. This shift could be relevant for areas receiving inputs from place cells at different points in the hippocampal dorso-ventral axis, which show a considerable increase in field size along the axis.

We showed that the phase-distance relationship is robustly present in area CA3. One possible caveat, also valid for the wild type mice analysis, is that since CA3 activity is very sparse, the analysis was only possible on data pooled over many recordings. It is still possible that the phase-distance distribution might show a different relationship for single session recordings with a high number of simultaneously active cells. One important difference compared to area CA1 is the presence of strong recurrent connectivity in area CA3, which could have an effect in the network coordination and the encoding of space at the population level, even if individual cells are theta modulated and exhibit phase precession. Further theoretical and experimental work should aim at better understanding the observed biased distribution of phases, the apparent uniformity of the down-phase group, and the possible implications these have for the encoding of spatial relationships.

The analysis also revealed that, in contrast to what we observed in wild-type mice, the phase-distance relationship was largely absent for Fmr1-null mice. One possible reason could be that individual place cells seem to be less organized by the phase of theta in the knockout mice than the wild type and the phase-frequency discharge probabilities of the place cells are also somewhat less coordinated in the knockout than the wild type (Talbot et al., 2018). It is interesting to compare the results that we found in Fmr1-null mice in Fig. 6 with that of the jittering analysis we performed in one of the rat datasets in Fig. 2D. In both the Fmr1-null mice and the rat data with sufficiently large levels of jittering the phase-distance relationship is lost. However, in the jittering analysis, as the bimodality of the correlation phase distribution disappears (at a *σ* value somewhere between 20ms and 25ms; see the polar plots at the top of Fig. 2D), the relationship between phase and spatial distance is also lost, as testified by the low value of *w* as *σ* increases. However, in the knock-out mice despite the disentanglement between the correlations phases and the place field distance, the distribution of correlation phases is far from uniform. One possible explanation for this could be that although the coordination of spikes between place cells are lowered in the knockout mice to a level that the phase-distance relationship is lost, there is still a significant enough level of theta modulation of the neurons that some structure in the distribution of phases remains. In any case, further work should address whether this phase code, relevant at behavioural timescales, is needed for long-term memory formation of independent representations of spatially informed tasks, since knock-out mice present deficiencies when required to perform discrimination tasks of past events with a spatial component, as opposed to performing as well as wild-type mice at learning and retaining unchanging spatial information (Radwan et al., 2016; Kooy et al., 1996; Krueger et al., 2011).

Besides the implications that the phase-distance relationship has on navigation (John O’Keefe and Burgess, 2005; Moser et al., 2008) and sequence coding and learning (Skaggs et al., 1996; Tsodyks et al., 1998; Melamed et al., 2004), one of us has previously shown that the relationship between correlations-phase and place field distance plays a crucial role in a model of the formation of grid cells (Monsalve-Mercado and Leibold, 2017). For phase coding to play this role, it needs to be valid during natural open field foraging, and to have an effect at timescales that are relevant for synaptic plasticity, which we have demonstrated in this paper. There are a number of ways downstream areas can be tuned to listen to the information encoded in the correlations on the theta channel. One possibility is that the architecture and dynamic properties of the receiving network favors responses to this frequency, as it is the case for some resonant circuits (Stark et al., 2013). Another important way is for single cells to directly listen to this theta channel. A relevant example are stellate cells in layer II of entorhinal cortex (LIIS), which exhibit inherent resonant behaviour at theta frequencies that is able to amplify the theta component of its input (Alonso and Klink, 1993; Giocomo et al., 2007; Haas and White, 2002). Moreover, since subthreshold resonance is equivalent to a band-pass filter of theta frequencies (Richardson et al., 2003), stellate cells could directly develop the hexagonal grid cell pattern through Hebbian learning by listening to the phase-distance relationship present in CA1. Principal cells in layer II of medial entorhinal cortex (mECII) do not receive direct input from hippocampal areas, but rather receive relayed input from layer Vb pyramidal cells (LVP), which likewise receive feedback input from LIIS cells (Witter et al., 2017; Fuchs et al., 2016; Nilssen et al., 2019; Buetfering et al., 2014). In addition, pyramidal cells in mECII (LIIP) also exhibit triangular patterns, and communicate with LIIS cells via intermediary cells. Filtering could be implemented there at the circuit level, with the entorhinal microcircuit LIIS-LVP possibly interacting with LIIP to generate grid patterns in all or a subset of cells within the circuit.

The theory makes an additional prediction for grid cells in the entorhinal cortex, which show strong phase precession in each of their firing fields (Hafting et al., 2008; Jeewajee et al., 2014; Climer et al., 2013). For idealized grid cells within a module, similar calculations as those reported in Monsalve-Mercado and Leibold, 2017 predict that averaging over all the firing fields of a grid cell pattern results in a bimodal distribution of phases similar to that of place cells in the Hippocampus. This is because each field effectively acts as an additional recording session. For grid cells belonging to the same module, a better measure of field overlap is provided by the spatial offset between the grid patterns. Previous work has taken steps in understanding the relationship between pairwise correlations between grid cells and their spatial phase differences. For instance, it has been shown that the absolute amount of spike-time noise-correlations decreases with the spatial phase difference between pairs of grid cells (Dunn, Mørreaunet, et al., 2015; Tocker et al., 2015), making pairs of grid cells with small spatial phase difference exhibit positive functional connections while those with larger grid spatial phase difference show negative functional connectivity (Dunn, Mørreaunet, et al., 2015). More recently it was shown that this relationship between pairwise correlations and spatial phase difference persists during sleep (Gardner et al., 2019; Trettel et al., 2019). Although Dunn et al. (Dunn, Mørreaunet, et al., 2015) did establish that the theta oscillation plays an important role in explaining the correlated activity of a population of grid cells, the relationship between spatial phase difference and functional connectivity did not qualitatively depend on phase precession. While these studies do shed light on the relationship between pairwise correlations and spatial phase difference in grid cells, none of them have focused on correlations at the theta time scale (theta band filtered correlations) to quantify the relationship between those correlations -which are highly influenced by phase precession- and the spatial relationship between grid cells’ firing patterns which would be the key prediction of the analytical results of Monsalve-Mercado and Leibold, 2017 extended to grid cells. It would thus be a natural next step to analyze recordings from mEC to see whether the phase coding demonstrated here for hippocampal place cells also exists for grid cells. This requires obtaining enough simultaneously recorded cells from the same module.

On the theoretical side, it would be interesting to extend the analysis reported in Monsalve-Mercado and Leibold, 2017 and relax some of the assumptions in the calculations, perhaps by relying on simulations. One may, for instance, study the relationship between the phase code and the size of the place fields, in particular in cases where the two place fields are different in size, the cells precess to different degrees, or where there are differences between the peak firing rates of the place cells, as normally happens in real data. Studying similar effects to this latter point would be particularly interesting to evaluate in grid cells where the variability in the peak firing rates across fields have been postulated to play a crucial role in the representation of spatial information (Dunn, Wennberg, et al., 2017; Kanter et al., 2017; Ismakov et al., 2017).

## Acknowledgements

The authors thank all experimental labs for their efforts in providing the datasets. M.M. is supported by the NSF NeuroNex Award DBI-1707398 and The Gatsby Charitable Foundation. M.M. and Y.R. were also supported by the Kavli Foundation, and the Center of Excellence scheme of the Research Council of Norway (Center for Neural Computation). M.M. thanks Christian Leibold for discussions in the early stages of the project.

